# Black chromatin is indispensable for accurate simulations of *Drosophila melanogaster* chromatin structure

**DOI:** 10.1101/2021.12.12.472204

**Authors:** Irina Tuszynska, Pawel Bednarz, Bartek Wilczynski

## Abstract

The interphase chromatin structure is extremely complex, precise and dynamic. Experimental methods can only show the frequency of interaction of the various parts of the chromatin. Therefore, it is extremely important to develop theoretical methods to predict the chromatin structure. In this publication, we describe the necessary factors for the effective modeling of the chromatin structure in *Drosophila melanogaster*. We also compared Monte Carlo with Molecular Dynamic methods. We showed that incorporating black, non-reactive chromatin is necessary for successfully prediction of chromatin structure, while the loop extrusion model or using Hi-C data as input are not essential for the basic structure reconstruction.

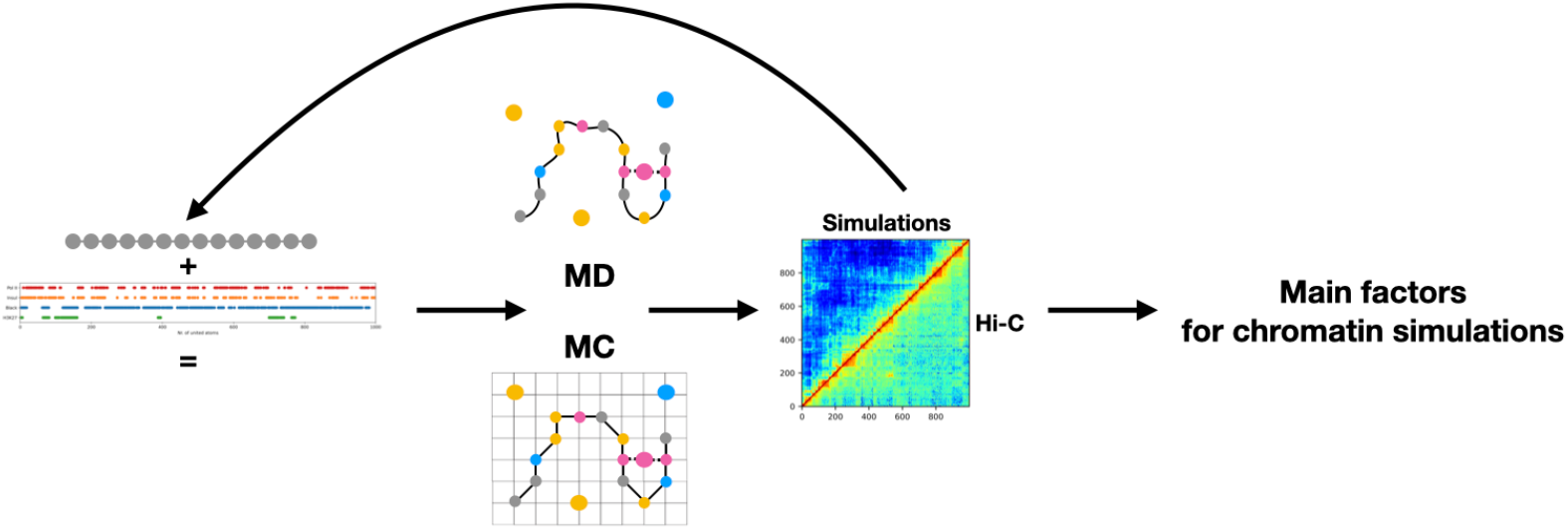

## Introduction

The nucleus of every living cell contains DNA that encodes all of the genes in an organism. It is estimated that each human cell contains approximately 2 meters of DNA polymer, folded into the cell’s nucleus that is 10 micrometers in diameter. Genomic DNA in the nucleus exists in the form of chromatin, which assumes complicated spatial structures necessary for the proper functioning of the cell in the interphase. The organisation of chromatin exists at various scales within the genome to provide different degrees of chromatin compaction [1]. Processes such as replication, DNA repair and transcriptional regulation require specific spatial structure of chromatin [2, 3], while for example silencing of chromosome X causes huge chromatin reorganisation [4]. Chromatin is a very dynamic creation and changes its conformation in response to the processes occurring in the cell nucleus. Abnormal structure of chromatin is linked with various diseases: ATR-X syndrome, ICF syndrome, Rett syndrome, Rubinstein–Taybi syndrome [5], cancer [6, 7]. Understanding the mechanisms that trigger chromatin folding, its patterns, and the factors they depend on, is a fundamental and still open question in modern biology that could help designing new personalised diagnostic tools for diseases connected with chromatin misfolding.

The spatial organisation of the genome is directed by intra- and inter-chromosomal interactions that involve components such as transcription factors, replication and transcription machineries, Polycomb bodies, insulators, cohesin as well as contacts with the lamina. Experimental methods like chromosome conformation capture (3C) and high-throughput derivatives like Hi-C do not allow to directly visualise the structure of chromatin, they show only the average frequency of contacts of individual regions of chromatin. Theoretical methods are extremely useful and effective tools to support experimental research. For example, methods based on Monte Carlo (MC) or Molecular Dynamics (MD) simulations were frequently and successfully used to reproduce the structure and dynamics of mobile biopolymers such as proteins or nucleic acids [8, 9, 10] as well as some chromatin properties [11].

Until now, a lot of successful attempts to predict chromatin or a part of chromatin structure were performed [12, 13, 14, 15, 16, 17]. In most of the work mentioned above, the authors model interactions between distant segments of the chromatin filament based on the contact frequencies determined by Hi-C. These experiments are therefore called data-driven simulations. For example MCMC5C program that uses data obtained by the methods 5C and Hi-C as well as Monte Carlo method utilize the Markov chain for modeling the structure of chromatin [12]. The Nicodemi group uses the Monte Carlo method with SBS (Strings and Binder Switch) model to simulate the main properties of chromatin [11] and MD with SBS model to predict the folding of a small part of chromatin [13]. The Marenduzzo group, in contrast to previously described models, used the fitting free model where information about polymerase, promotors, enhancers and HP1*α* binding sites with MD method simulations were used to obtain chromatin structure [15]. Authors compared the generated contact maps with Hi-C data by measuring the similarity of the TADs boundaries, which were determined by visual inspection, leading to an imperfect measurement of similarity between the original and generated contact maps. While reconstructing the TADs bounders is a useful step towards judging the results of simulations, however more quanitative are needed for TADs determination and computing contact map similarity to be able to assess which modeling approach is better than the other. Another model of chromatin folding is related to the active loop-extrusion mechanism where cohesins forms extrusion loops, while the CTCF serves as an anchor for the cohesin to stop the extrusion process [18]. While ShNeigh method combines the MDS technique with local dependence of neighboring loci modeled by a Gaussian formula, to infer the best 3D structure from incomplete contact frequency matrices [16]. However until now it is still unclear which factors drive an effective simulation of chromatin structure.

To address this issue, here we present an *in silico* investigation of the main chromatin factors important for the simulation of chromatin structure on the example of 2L chromosome of *Drosophila melanogaster*. Instead of using the information obtained by experimentally-determined Hi-C data to guide the simulation, we start with 1D features that describe chromatin epigenetic state including insulator proteins, polymerase and histone modifications and were obtained independently from the Hi-C data. Then we use these inputs to set up a simulation and generate a population of possible chromosome structures. Only then, we compute the average contacts from multiple simulations and compare the resulting contacts with those seen experimentally in Hi-C. Due to the fact, that the evidence of the existence of active motor proteins that pull chromatin to specific locations of the nucleus is currently very debatable, we are not using any additional forces to pull contacting fragments of chromatin to each other, even relatively short range Lennard-Jones potential. We just applied a simple SBS model with Metropolis Monte Carlo algorithm without active extrusion mechanism of chromatin. We also compared the Monte Carlo method with Molecular Dynamics for chromatin structure simulations.

## Methods

### Implementations of Monte Carlo simulations

To simulate the chromatin structure, we implemented a procedure based on the one described by Barbieri et al. [11] and created the ChroMC program. Our implementation is freely available (https://github.com/regulomics/chroMC) to other users, while the prototype described by Barbieri et al., as far as we know, is not publicly available. In our program, besides the Monte Carlo chromatin simulation described by Barbieri et al., we also implemented lamina-like binders to possibly take advantage of the interactions of floating proteins with the nucleus lamina in the simulations of the chromatin structure. Our ChroMC uses the strings and binders switch (SBS) polymer model where chromatin is represented as a polymer chain of connected united atoms. Some of chromatin united atoms have discrete binding sites for freely diffusing atoms that represent proteins interacted with chromatin. All atoms are placed in the 3D grid. The polymer has self-avoiding-walk properties, and all united atoms move in space by diffusion, simply under the constraint of no overlap. We implemented Metropolis Monte Carlo algorithm. The scoring, energy-like function increases, when the corresponding diffused atom is at the distance below a certain threshold to the corresponding chromatin binding site. In each step of the simulation, if the energy increases, the new structure is accepted. In the opposite case, when the scoring function decreases, the new structure is accepted with probability proportional to exp (−ΔH/kBT), where ΔH means the score difference between the previous and the new step of the simulation. One binding site could bind 6 proteins that could be of the same or different types, which also differ our implementation from the Barbieri et al. model, where each chromatin bead can interact with one specific binder. Our implementation describes the natural environment of chromatin, where complexes of many proteins are bind to chromatin in particular range of nucleotide (one bead of chromatin in simulation corresponds to 5 Kbp). We should also remember that one bead of our simulation represents several thousand base pairs.

### Monte Carlo simulations

We run all simulations on the first 1000 united atoms of the 2L *Drosophila melanogaster* chromosome due to the limited processor time. One united atom represents 5 Kbp of chromatin. We used Hi-C maps with the same resolution to check the results of our simulations. Hi-C maps were kindly provided by the Furlong laboratory. We utilized ChIP-seq data to define binding sites of appropriate proteins on chromatin. All ChIP-seq data with the exception of five colors of chromatin (GSE22069 [19]) and the Nipped protein data (GSE118484 [20]) was taken from the Modencode database [21]. All factors that can influence the structure of chromatin and which were used to simulate it are listed in Supplementary Table S1.

The values in the contact map are equal to 1/d, where d means distance between united atoms.

### Calculation of contact map and Hi-C map similarity

To quantify the similarity of the contact map with the Hi-C map we used two methods: global similarity and local similarity calculation.

To find global similarity, we determine the Spearman correlation coefficient for each diagonals separately (correlations for the first diagonals, second diagonals etc., starting from the main diagonal of the map) and then we calculated the average value for the specific range of diagonals (first 100, 100-200, 200-300 or the first 300). We considered correlation for the first 100 diagonals as the main measure of similarity since most contacts are in the first 100 diagonals (Figure 1A black line). Each point on the global correlation plot represents the correlation between the individual diagonals (Figure 1B).

**Figure 1:**
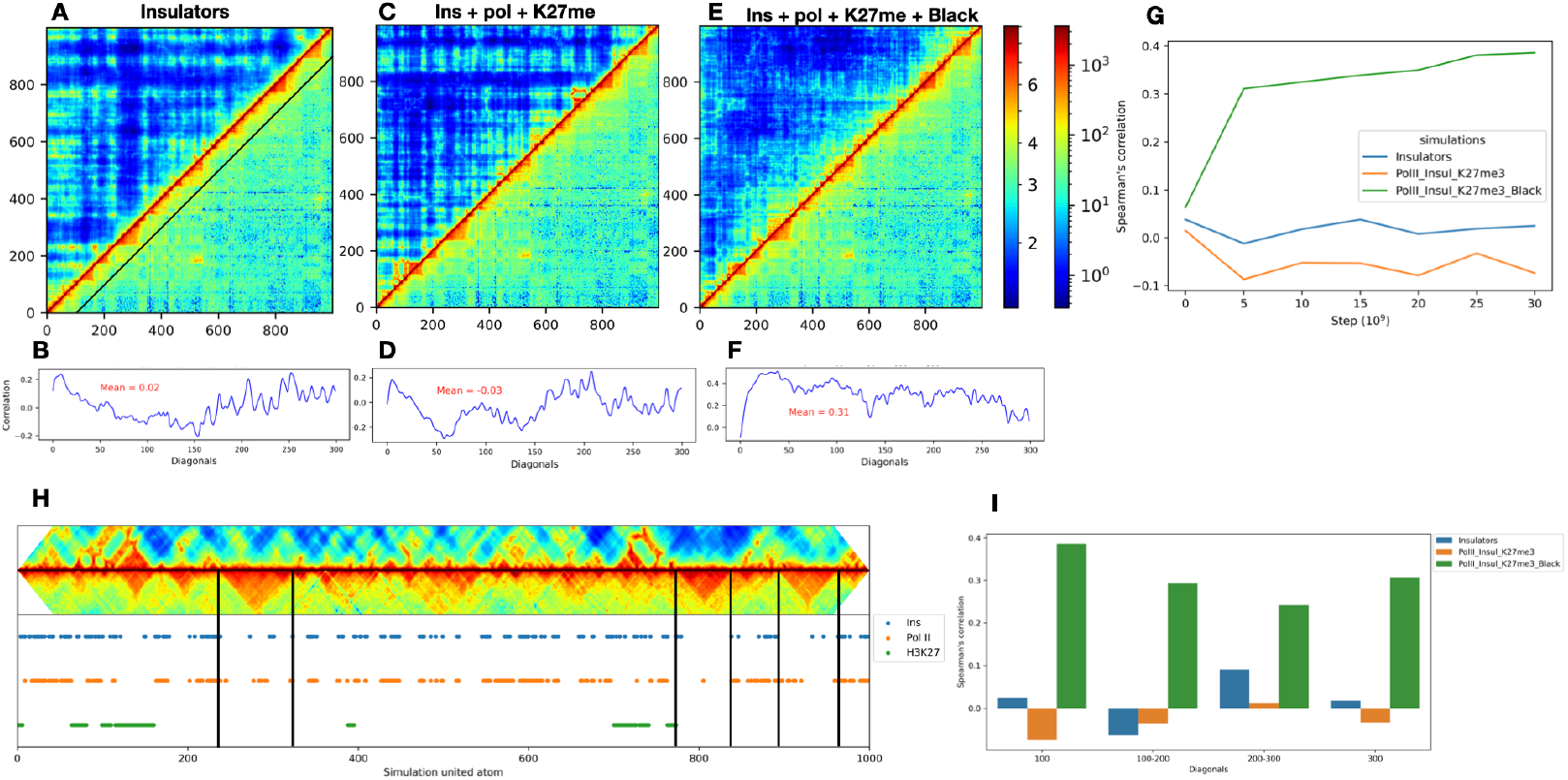
A successful simulation pathway of the first 5 Mbp *Drosophila* 2L chromosome. A: Comparison of the contact map built for the last step of twenty independently run of the Monte Carlo simulations with only information about insulators binding sites (upper left triangle) with 5 Kbp resolution Hi-C map (lower right triangle). B: The plot shows the Spearman correlation coefficient for each diagonal between the generated contact map and the Hi-C map. C, D and E, F show the same graphical representation of the similarity of the contact maps with the Hi-C map. Contact maps were generated for simulations with information about insulators, polymerase II and H3K27me (C, D) and for simulations where information about insulators, polymerase II, H3K27me and black chromatin (E, F) was included. G: Changes of correlation over time for three mentioned simulated conditions. The contact map was calculated for 20 trajectories. H: Positions of chromatin interaction sites with considered proteins with respect to Hi-C domains. I: Correlation coefficients for the contact maps created for the last step of simulations, calculated for different diagonal number thresholds.

For local similarity, we calculated the average correlation for each diagonal in a small window of a certain size (100 × 100 united atoms in our case). Next the window is moved across the diagonal by the defined step (15 united atoms in our case). Each point on the local similarity figure represents the mean correlation for all diagonals in the small window (Figure 1 B, D, F).

### Molecular Dynamic simulations

We used the LAMMPS program [22] for Langevin dynamics simulations of a similar model to that used in Monte Carlo simulations. Chromatin fibers are represented as a coarse-grained model, where beads representing a given number of base pairs (5 Kbp in our case) are connected by bonds simulated as a spring using FENE potential (maximum extension 1.6 times bead diameter). The polymer is given flexural stiffness by a cosine interaction between the triangles of adjacent beads. Protein - protein and chromatin bead - chromatin bead interactions only consist of a repulsion component using a Weeks– Chandler–Anderson potential. Interactions between proteins and their binding sites on chromatin fiber are represented by Lennard–Jones potential that has short-range repulsive and longer-range attractive parts. All beads are confined within a box with periodic boundary conditions. We utilize the same numbers of binders and binding sites as were used with the Monte Carlo method. We used the same configuration files as was described in the Marenduzo publication [14] and changed them slightly to represent our models of chromatin. We performed from 5 to 39 repeats of simulations for different sets of factors influencing the chromatin structure.

## Results

### Searching for a set of factors that interact with chromatin needed to simulate the chromatin structure

In order to find the minimal set of proteins interacting with chromatin that are required to simulate the chromatin structure accurately without any prior knowledge from Hi-C map, we tested different combinations of available proteins (Table S1).

As it is well established in literature that insulator proteins (in the case of *Drosophila*, we count BEAF, CTCF, CP190 to that class) can have a decisive influence on the chromatin structure as they are found with other border-associated proteins at domain boundaries [23], we have begun by using Monte Carlo simulations with data on insulator binding sites only. Some of *Drosophila* insulator proteins are known to interact with lamin (SuHW and CP190) [24]. We added the lamin layer to our model to take into account chromatin interactions with the lamin. In order to understand if the simulations are close to the structure of chromatin established by Hi-C, we construct the summary contact map of the last steps of multiple simulations running independently and compare the contact map with the Hi-C map. The contact map was prepared for 20 structures, each of them was the last frame of MC simulation 30 ×10^9^ MC steps long. However the contact map looks very different comparing to a real Hi-C map for 12-14 h after fertilization (Figure 1A). To numerically compare the contact map that summarised our simulation with the Hi-C map, we calculated the Spearman correlation coefficient for the first 100 diagonals as the most frequent contacts appear until around the first 100 diagonals (see black line on the Figure 1A). Next we calculated a mean Spearman correlation for the first 100, 100-200, 200-300 and 300 diagonals and found that there is no correlation between contact map obtained after MC simulations with information about insulators and Hi-C map (Figure 1A). The correlation does not exceed 0.1 value throughout the whole simulation for the first 100 diagonals and for all diagonal thresholds (Figure 1A and 1I). We also could observe that correlation coefficient has no tendency to growing up during the simulation (Figure 1G).

Next we checked if the addition of data on active and inactive chromatin is enough to simulate chromatin structure, as it was done by Brackley et al. [15]. As an active chromatin we used polimerase II binding sites, while histone modification K27M3 represents a closed heterochromatin. A contact map of the simulated chromatin still looked very different to the real Hi-C map (Figure 1C), although locally one can observe improvement in some regions of simulated chromatin. TADs between 120 and 180 united atoms as well as between 700 and 750 united atoms on Figure 1C were well represented using information about insulators, polymerase II binding sites and H3K27 methylation.

At this stage of our project we tested simulations with and without lamina and unfortunately, contacts with lamina had a slowing effect on our simulations without much observable change to the resulting contact map (Figure S1). Based on these observations we decided to run all simulations without the lamin binding sites.

We also checked the effect of Nipped, which is the Drosophila derivative of Nipbl, on the simulation of the chromatin structure, as this protein recruits cohesin and is involved in the loop extrusion process [25], but adding NippdB interaction sites to the model did not bring any significant effect (Supplementary Table 1).

When we compared the results of simulations, where insulators, active and inactive chromatin were taken into account, we observed that the largest domains observable in the Hi-C map have very few contacts in our simulations (Figure 1H). This observation indicated, that some key component that was affecting the domains observed in Hi-C data was still missing in our model. In the publication of Filion et al. [19] authors found out that the largest domains usually belong to so-called “black” inactive chromatin, with the name coming from the lack of most binding signals in the DAM-ID experiments. According to their observations, the black chromatin can be described mainly by five proteins: H1, D1, IAL, SUUR, and LAM. We defined united atoms that represent binding sites for black chromatin proteins if at least four out of five mentioned above proteins are identified as bounded to the nucleotide range of the particular united atom. It should be noted that Filion et all. prepared location maps for five colors of chromatin on the Kc167 cell line, while we compared the simulation results with Hi-C map for embryos 12-14 hours after fertilization. Despite this the contact maps is much more similar to the Hi-C map (Figures 1 E, F) with correlation for the first hundred diagonals increased to 0.39 from nearly 0 in the previous cases (Figure 1G).

### An appropriate representation of the black chromatin

After we found out the parameter set that could reproduce the Hi-C map, we increased the number of trajectories to 39 which slightly improved the correlation (for 100 diagonals from 0.39 to 0.41). All next contact maps were presented for 39 trajectories.

To understand the best way of definition of the black chromatin, we compare our results with simulations, where the black chromatin sites are taken from the Filion et al. publication as well as where histone H1 interaction sites were used as a representation of the black chromatin, as, based on publication [19], H1 describes black chromatin localization with 80% overlap. We also run the simulations, where each color of chromatin that was defined by Filion et al., was considered as a separate type of interaction sites on the chromatin filament. However the best results were obtained for simulations with our definition of the black chromatin, where only chromatin united atoms are considered as the black chromatin if four out of five proteins were bonded (Figures 2A and S2). The greatest advantage of our definition of black chromatin is noticeable in the first hundred diagonals. However, even addition of H1 binding sites to simulations with insulators, polymerase II and H3K27me increase similarity of contact map and Hi-C from from -0.05 to 0.22 (Figure S2). It is clearly seen that both correlation for each diagonal profile (Figure 2B and 2F) and local window correlation (Figure 2D and 2H) are higher when we compare simulation with H1 (insulators + polymerase II + H3K27me3 + H1) with successful set of chromatin interaction compounds (insulators + polymerase II + H3K27me3 + black chromatin) (Figures 2B, C, D) than if we compare H1 simulation with those compounds without black chromatin (Figures 2F, G, H). Simulation with H1 to some extent reconstitutes large domains in the range of about 240-310, 790-840 and 900-965 beads, that are not visible in the simulation without the information about black chromatin (Figure 2C, G).

**Figure 2:**
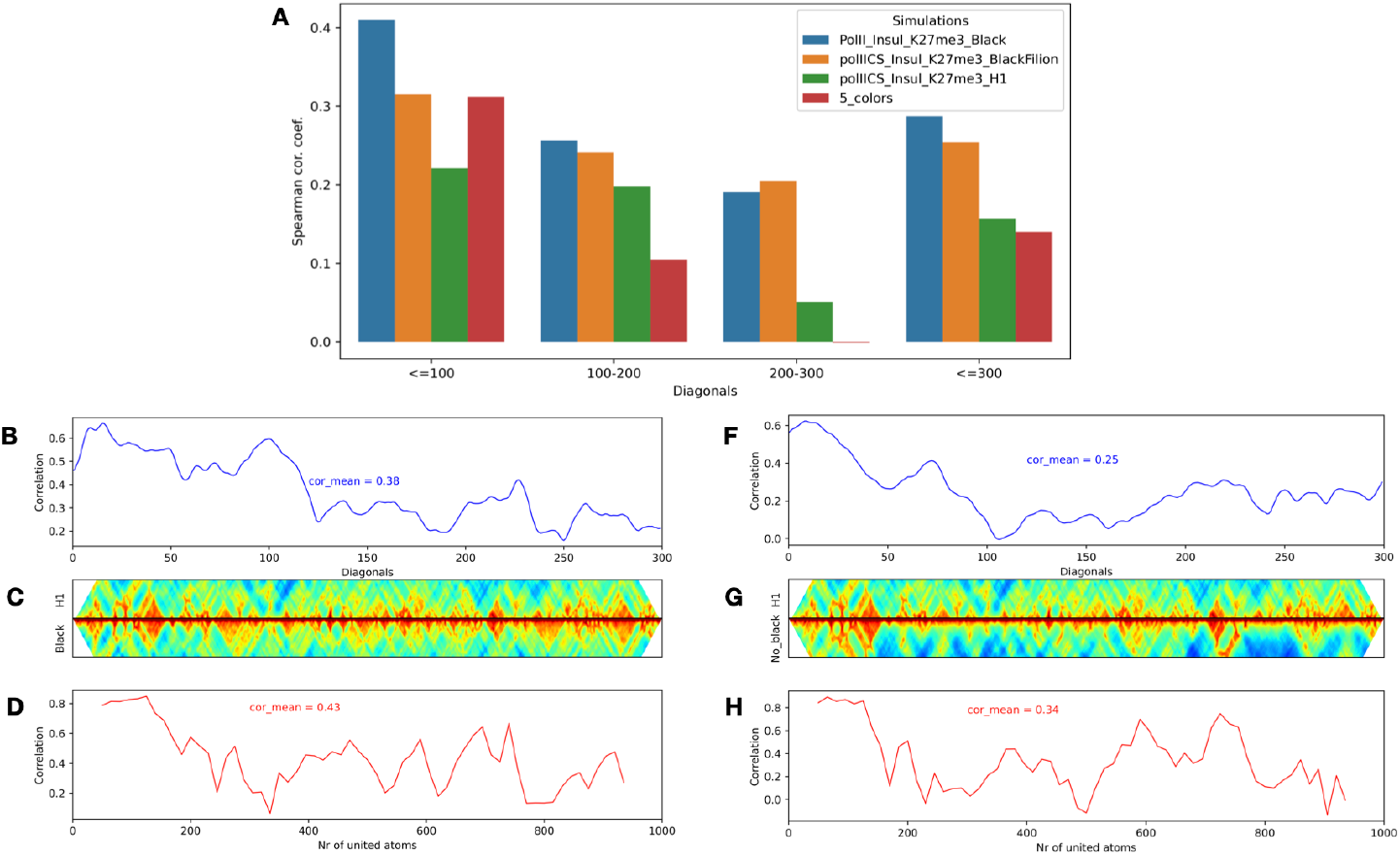
Different ways of defining black chromatin. A: Correlation for different diagonal thresholds when comparing the contact map with the Hi-C map for simulation with distinct definitions of black chromatin. B, C, D: Comparison of the simulation with the entire set of chromatin factors that were included in this study, with the same set, but for the definition of black chromatin, instead of the presence of four out of five proteins, histone H1 binding sites were used. F, G, H: Comparison of simulations without black chromatin with simulations with H1 interactions site as a representation of black chromatin. B and F - correlation coefficient as the function of individual pairs of diagonals. D and H - local correlation plot which is informative in combination with the contact maps C and G.

### Searching for the minimal set of factors that allow to simulate the chromatin structure

Since the insulator protein locations alone do not constitute sufficient input to the simulations leading to accurate chromatin contact maps (Figure 1A), we decided to eliminate the insulator binding sites from the chromatin binding protein configuration that can allow for successful recreation of chromatin contacts. In the same way, we checked the effect of the removal of polymerase and the modification of K27me3 on simulations. The correlation calculated for diagonals of the summary contact map from the last step of 39 simulation and the Hi-C map shows that removing any of the factors mentioned here (Insulators, PolII, H3K27me3) slightly reduces the correlation between simulation and Hi-C (Supplementary Tables S2, Figure 3). However, it appears that PolII is particularly important for interactions longer than 100 unified atoms of the chromatin filament, while insulators are most relevant for interactions within up to 100 unified atoms (Figure 3A). After calculating the local similarity for the simulations without H3K27me, a clear decrease in correlation can be observed in the region around the position 700, where this heterochromatin marker occurs. In the absence of polymerase, the greatest decrease, almost the same as in the absence of black chromatin, can be observed in the region around positions 180 - 200 (Figure 3B).

**Figure 3:**
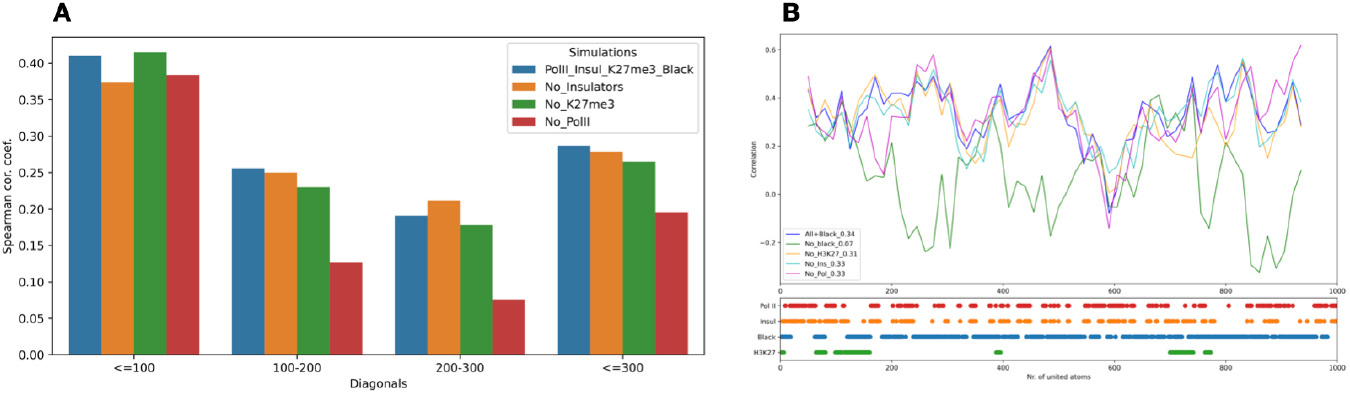
Selection of the smallest set of factors influencing chromatin structure simulation. A: Correlation of the Hi-C map with the contact map built from the last structure of 39 simulations. Correlation was calculated for different diagonal thresholds of the contact map. B: Comparison of the local correlation between the simulations with all elements causing successful chromatin simulations (insulators + PolII + H3K27me3 + black) with simulations without each of the above-mentioned factors affecting the chromatin structure.

### Comparison Monte Carlo with Molecular Dynamic simulations

We also checked if Molecular Dynamics (MD) will give the similar results for the parameters that were used for Monte Carlo (MC) simulations.

We run MD simulations with the set of model components(insulators, polymerase II, H3K27me marker and black chromatin locations) that produce contact map most similar to the actual Hi-C map. The mean correlations for the first 100 diagonals of MC and MD contact maps with Hi-C map are very similar (0.39 and 0.45 respectively) (Figures 4A, B and C, D), as well as similarity MC and MD to each other for the first 100 diagonals is rather high (correlation 0.51) (Figure 4E, F). For the longer chromatin interactions the correlation for MD simulations contact map with Hi-C sharply decay compared to MC contact map (Figure 4G).

**Figure 4:**
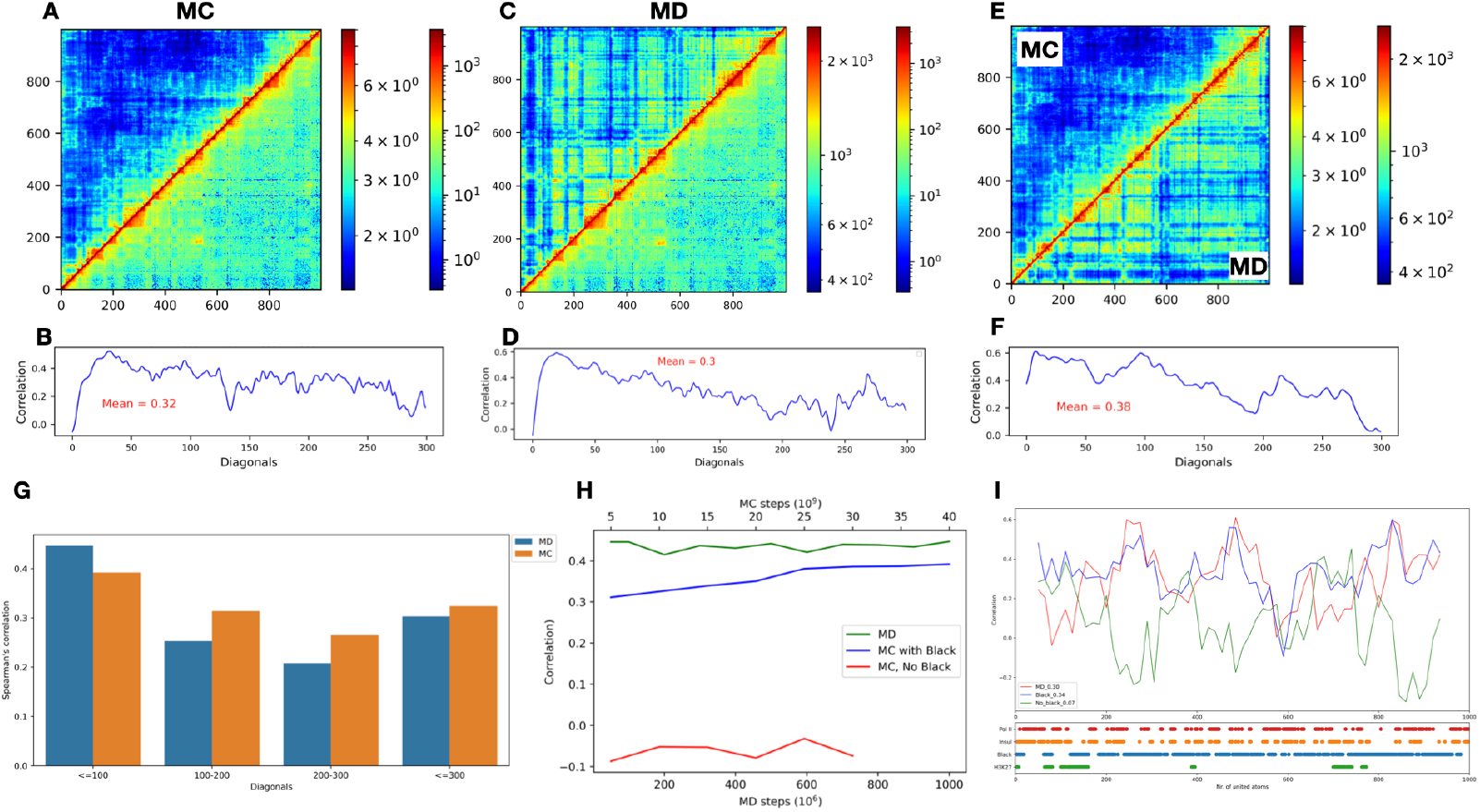
Comparison of Molecular Dynamics and Monte Carlo simulations. A and C: The contact map of MC and MD simulations respectively, with all chromatin factors that cause successful simulation. Top left triangle represents contact maps of the simulated structure, while bottom right triangle - the Hi-C map. B, D: plots represent the correlation between simulated contact map (MC and MD respectively) and Hi-C map for the first 300 diagonals. E and F: The same comparison between contact maps built on the last structure of 20 MC simulations and 20 MD simulations. G: Correlation for simulated MC and MD maps with Hi-C map for different diagonal ranges. H: Correlation changes over time for the first 100 diagonals for MC and MD simulations with all chromatin factors necessary for a successful simulation and MC without black chromatin information. I: Top - Local correlation for MC and MD with all chromatin factors (also black chromatin) and MC without black chromatin. Bottom - binding sites for all chromatin factors that affect the chromatin structure in simulations and were used in MC and MD simulations.

Both simulation methods give similar results for the first 100 diagonals, however MD simulations were relatively faster to achieve relatively high correlation of the first 100 diagonals (Figure 4G), while the longer range interactions seems to be less accurately represented (Figure 4G). The comparison of the correlation coefficient for the first 100 diagonal shows the advantage of the Molecular dynamics simulation over Monte Carlo (0.44 and 0.39 respectively) (Figure 4H), however, if we compare the mean of all local correlation coefficients, MC slightly outperforms of MD (0.34 and 0.30 respectively) (Figure 4I). An interesting observation can be made that in the area, where both the entire set of used factors and simulations without black chromatin have very similar correlation with Hi-C map, while MD method has a clear decrease in correlation (Figure 4I, around 80 - 130 united atoms and 680 - 750 united atoms). Both areas are at the position where the H3K27 marker is located which makes it possible for more types of binder atoms to bind. However, the number of bonded atoms in MC was limited to 6 and additionally limited by the mesh on which the simulated atoms reside. While in MD atoms were placed in the continuous space, where the limitations occur only due to the forces describing the unified atoms and the geometry of the atoms. These facts could make it more difficult to predict an appropriate structure of chromatin in the places where more than 3 types of united atoms could bind.

Comparing simulations made by ChroMC and LAMMPS both applications have their own strengths and weaknesses (Table 1). LAMMPS is a more powerful application, with possibility to run parallel processes, causing fast calculation, however it is really not easy to run stable MD simulation using LAMMPS for the starting user, while ChroMC can run even a user without experience in the area of structural bioinformatics.

**Table 1:**
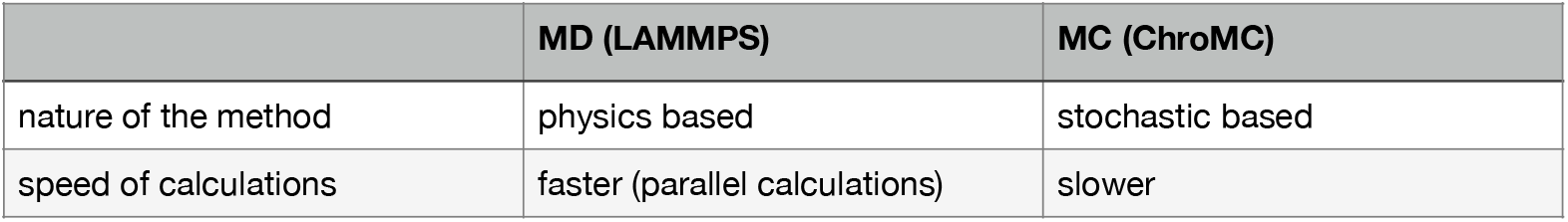
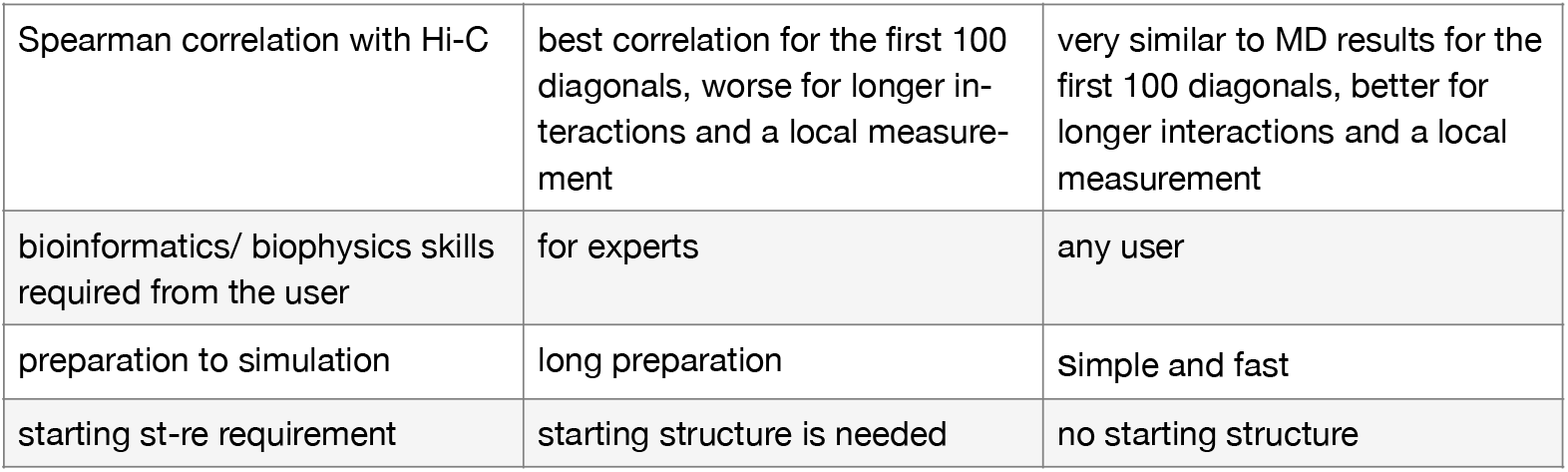
Comparison of MC and MD methods

|  | **MD (LAMMPS)** | **MC (ChroMC)** |
| --- | --- | --- |
| nature of the method | physics based | stochastic based |
| speed of calculations | faster (parallel calculations) | slower |
| Spearman correlation with Hi-C | best correlation for the first 100 diagonals, worse for longer interactions and a local measurement | very similar to MD results for the first 100 diagonals, better for longer interactions and a local measurement |
| bioinformatics/ biophysics skills required from the user | for experts | any user |
| preparation to simulation | long preparation | Simple and fast |
| starting st-re requirement | starting structure is needed | no starting structure |

## Discussion

To find the best minimal set of the chromatin factors that allow to predict chromatin structure we used different combinations of many components (Table 1S). Some of them, such as interactions with Nipped protein, related to the loading of cohesin into chromatin, showed no effect on the chromatin structure. CTCF and other insulators also showed very little effect on simulation of chromatin structure that could be seen only after removing insulator interactions sites from the successful simulations (Figure 3A, B). The interactions with lamin (SuHW and CP190), modelled as a separate chromatin interacting agent, also does not seem to be significant for the simulation of the chromatin structure (Figure S1).

Based on our experience the best set of components needed for the simulation of *Drosophila* chromatin consist from K3H27me marker, insulators (BEAF, CTCF, CP190) and polymerase II binding sites as well as the location of the black chromatin defined as the presence of at least four out of five binding sites of proteins: H1, D1, IAL, SUUR, and LAM. However, it is much more demanding and expensive to perform five experiments to obtain the interaction sites of five mentioned proteins. Fortunately for the sufficient description of the black chromatin even addition the information about H1 to the simulation could produce satisfying contact maps of chromatin. The main role in the formation of an appropriate signature of contacts that is seen on the Hi-C map has the black chromatin. As was described previously, our simulations show that the black chromatin is responsible for formatting the biggest domains of chromatin, as described by Filion et al.. Whereas polymerase II and H3K27me3 are needed for maintaining a few not very large domains. We can see that excluding any component from the list: insulators, polymerase II and H3K27me3 decrease similarity with Hi-C map only slightly, while lack of black chromatin cause the contact map almost completely loses its resemblance to the Hi-C map. The lack of noticeable effect on the chromatin structure of removing CTCF and other insulators in the simulations is in agreement with studies where knock-outs of CTCF had only minor effects on domain organization [26, 27], which again suggests that this factor cannot be the main organizer.

The key role of black chromatin in simulating the chromatin structure could be investigated in other higher eukaryotes. It might be the case, that using a real lamin binding would be equally beneficial for the simulations, however in our experience they were slowing the simulations down too much to see their potential benefit.

We show that both Monte Carlo and Molecular Dynamic simulations produce similar results and are able to simulate the chromatin structure with resolution of 5 Kbp with the help of a simple model of chromatin dynamics without taking into account the loop extrusion mechanism. The main advantage of ChroMC comparing with LAMMPS is speed and ease of setting up the simulation.

In summary, simulations involving only one-dimensional information corresponding to different chromatin states provide a robust, simple and generic mechanism that can concentrate specific proteins associated with related sites into clusters and fold the chromatin at the interphase in a manner known in vivo. The main influence on the appropriate chromatin structure has black chromatin. Speaking of simulation methods, both MC and MD methods can be used to simulate chromatin, while the ChroMC method developed by us is freely available and easy to use even for users inexperienced in molecular simulations.

## Data Availability

The code is available at https://github.com/regulomics/chroMC

## Acknowledgements

This work was supported by the National Science Centre grants [2014/15/N/NZ2/00397] and [2015/16/T/NZ2/00178] to PB and [2015/16/W/NZ2/00314] to BW and IT

## Author Contributions

All authors contributed to conceptualization of the study. PB wrote a prototype implementation of the ChroMC, IT improved the software, performed all experiments, analyzed the results and wrote the first version of the manuscript, that was then revised by BW and IT. All authors read and approved the final version of the manuscript.

