## Supplemental Figure S1, S1 and table S1 for "Black chromatin is indispensable for accurate simulations of *Drosophila melanogaster* chromatin structure"

We run simulations with polimerase II binding sites and H3K27me3 data both with lamin (SuHW and CP190) and without lamin interactions. Next, we compared contact maps created for 5 trajectories after 40 * 10**9 Monte Carlo steps with Hi-C map. The lamin does not improve the contact map similarity to Hi-C map (Figure S1 A and B). The mean Spearman’s correlation for trajectories with lamin with Hi-C map for the first 100 diagonals was -0.06, while for trajectories without lamin the mean correlation for the first 100 diagonals is -0.01. The similarity between contact maps for trajectories with and without lamin is easy observable and measured ( correlation for the first 100 diagonals is 0.4), however the similarity growing to 0.52 when we created the contact map from 10 trajectories with lamin and compared it with contact map created from 5 trajectories without lamin (Figure S1 C,D,E).

Figure S1: Similarity between simulations with lamin binding sites and without lamin binding sites. A and B: Top - a heatmap with the Hi-C map (lower right triangle) and the contact map (top left triangle) prepared using the last step of 5 simulations with polymerase II, H3K27me3 and lamin (SuHW, Cp190) informations (A) or polymerase II and H3K27me3 (B). Bottom - The correlation of each diagonals in Hi-C and contact map showed on the top heatmap.  C: Top - a heatmap with the contact map for simulations without lamin (lower right triangle) and the contact map for simulations with lamin (top left triangle). Bottom - correlation for the first 300 diagonals of showed heat map. D: Comparison of the heat map for 10 trajectories with lamin with the contact map of 5 trajectories without lamin. Bottom - correlation for the first 300 diagonals. E: Comparison of the contact map for 5 trajectories with 10 trajectories with lamin. Botto - the plot of the correlation as a function of the first 300 diagonals.

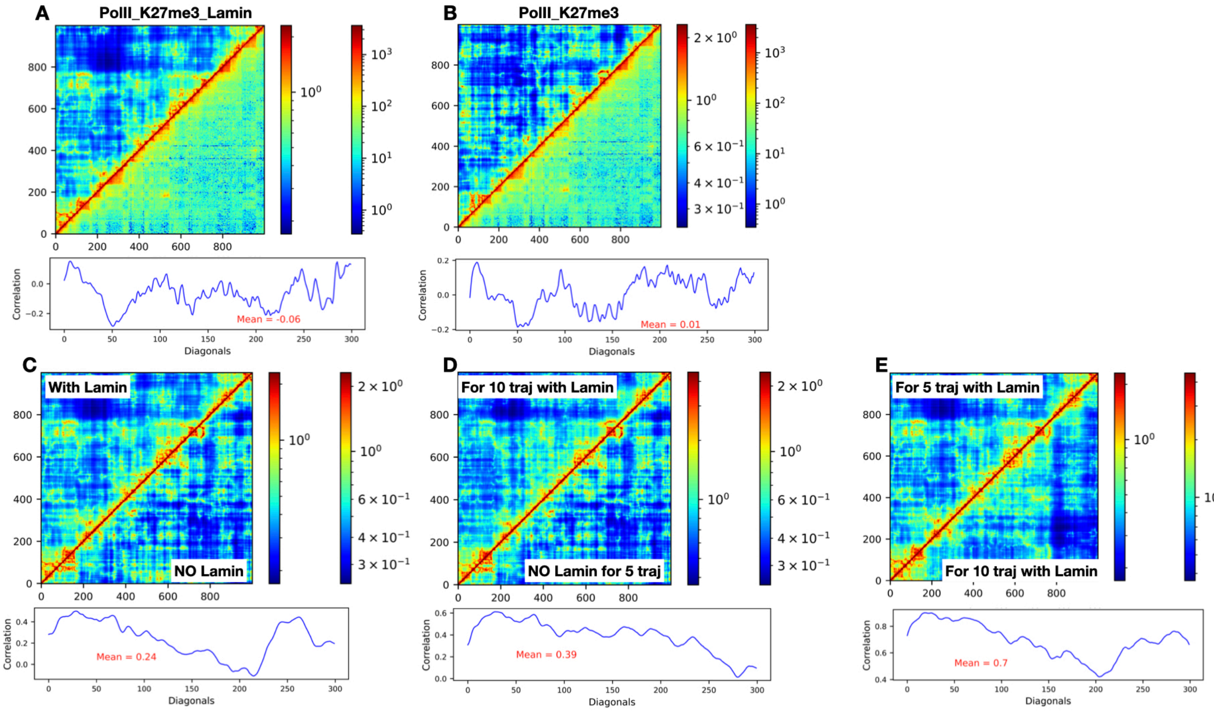

Figure S2: Comparison of different methods to define black chromatin. A, B, C: top left triangle - simulations with black defined as presence at least 4 out of 5 proteins, H1 binding sites or defined in Filion et al. respectively. Bottom right triangle - hi-C matrix, D: 5 colours of chromatin in definition from FIlion et al. was used for running of the simulation and compared with Hi-C map, E: simulation with insulators + polymerase + H3K27me + H1 as a definition of black chromatin is compared with insulators + polymerase + H3K27me + black defined as 4 out of 5 proteins. F: top left triangle the same as on E, but compared with simulation without black chromatin.

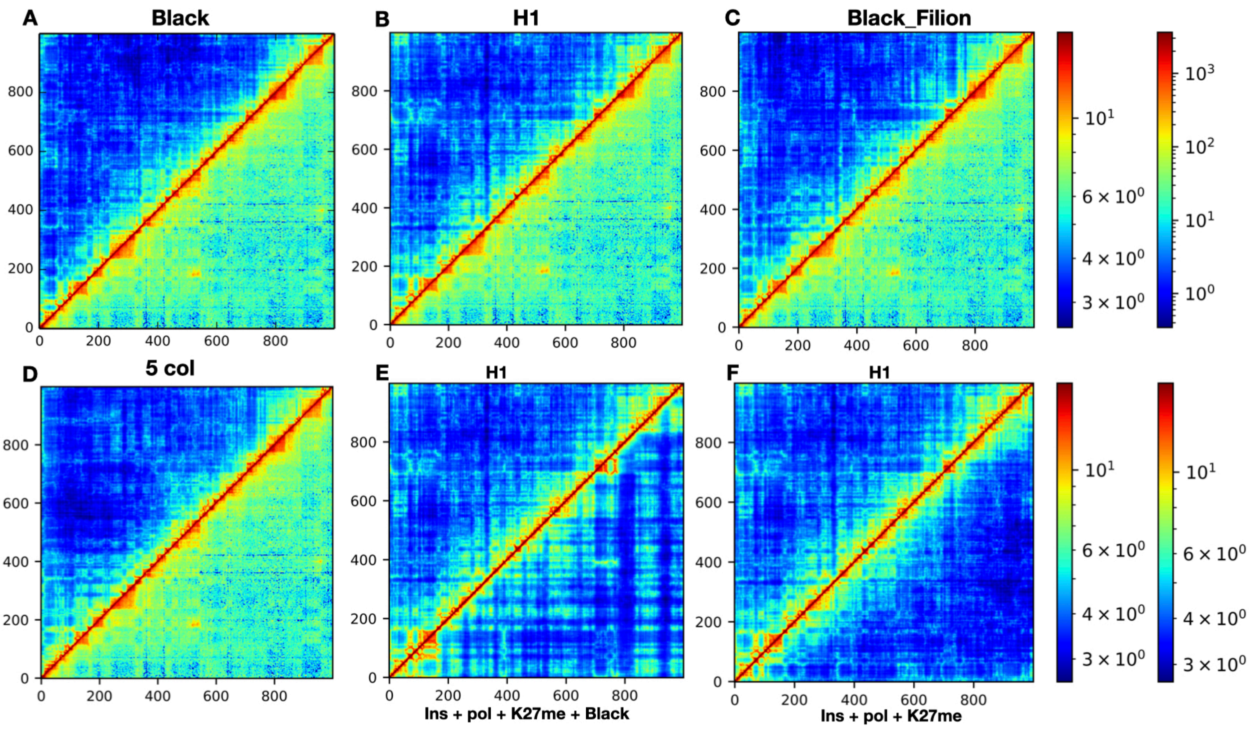


Table S1: Factors that were taken into consideration in the various simulations and the highest correlation for the first 100 diagonals of the contact map and the Hi-C map for each simulation. The contact map was made using the last saved structures from 14 simulations. * - contact map was built based on the structures from 5 simulations. The highest correlation is bolded.

Chromatin factors used in simulations

| MC Simultions | Insulators (BEAF. CTCF. CP190) | Lamin (SuHw. CP190) | Nipped | Polimeraze II | K27me3 | Black: 4 from 5 proteins | Black: from Filion | 5 colours | H1 | The best correlation for 14 simulation |
|---|---|---|---|---|---|---|---|---|---|---|
| Ins_Lam | + | + | | | | | | | | 0.02 |
| Pol_K27me3 | | | | + | + | | | | | -0.04 |
| Ins_Pol_K27me3 | + | | | + | + | | | | | 0.03/ -0.05* |
| Ins_Lam_Pol_K27me3 | + | + | | + | + | | | | | -0.01* |
| Ins_Pol_K27me3_BlackFilion | + | | | + | + | | + | | | 0.3 |
| Ins_Pol_K27me3_Black | + | | | + | + | + | | | | **0.41** |
| Ins_Pol_K27me3_H1 | + | | | + | + | | | | + | 0.19 |
| 5 colors | | | | | | | | + | | 0.28 |
| Ins_Pol_K27me3_Nipped | + | | + | + | + | | | | | -0.08 |
| Pol_K27M3_Black | | | | + | + | + | | | | 0.37 |
| Ins_Pol_Black | + | | | + | | + | | | | 0.37 |
| Ins_K27M3_Black | + | | | | + | + | | | | 0.34 |
